# Synchronised brood transport by ants occurs without communication

**DOI:** 10.1101/364273

**Authors:** Danielle P. Mersch, Jean-Pierre Eckmann, Alessandro Crespi, Laurent Keller

## Abstract

Collective behaviours in societies such as those formed by ants are thought to be the result of distributed mechanisms of information processing and direct decision-making by well-informed individuals, but their relative importance remains unclear. Here we tracked all ants and brood movements to investigate the decision strategy underlying brood transport in nests of the ant *Camponotus fellah*. Changes in environmental conditions induced workers to quickly transport the brood to a preferred location. Only a minority of the workers, mainly nurses, participated in this task. Using a large number of statistical tests we could further show that these transporters omitted to recruit help, and relied only on private information rather than information obtained from other workers. This reveals that synchronised group behaviour, often suggestive of coordinated actions among workers, can also occur in the complete absence of communication.

## Introduction

The success of group actions frequently relies on communication between individuals. Communication is manifest in animal groups as different as jellyfish that use bioluminescence to locate each other and team up^1^, prairie dogs that call to warn their family of predators^2^ and honeybees that use waggle dance to signal a food source to nest mates^3,4^. In all these cases communication serves to enhance the efficiency and safety of the group. However, communication is complex. It requires that the sender recognizes the appropriate circumstances and produces a correct signal, and that the receivers are able to understand the signal and react appropriately. These inherent difficulties constrain when and under what conditions groups of animals might communicate.

In ant societies communication is widespread and individuals make use of an array of olfactory, vibrational and tactile communication strategies. Therefore, communication is often assumed to be underlying all group behaviours^5,6,7,8,9,10^. Ants optimize foraging by creating pheromone trails^11,12^, and by recruiting help to retrieve food through tandem runs, a method whereby a knowledgeable ant induces a naive ant through tactile and chemical signals to follow it^13^. In emergencies, ants release highly volatile alarm pheromones^11^. If a nest is destroyed knowledgeable ants first lead tandem runs to new nest sites before switching to brood transport^14^. In all these instances communication is manifest and beneficial to the society. Pheromone trails and tandem recruitment reduce the risks of random food searches and ensure that a sufficient number of workers locate and retrieve food before it disappears, thereby enhancing the colony’s chances of survival and reproduction. Similarly in emergencies the survival of the colony is at stake. Alarm pheromones ensure that workers are alerted and leave the nest^15^ for fight or flight. Tandem runs ensure that a sufficient number of workers know the location of a safe alternative nest before evacuating brood^9^. However, there is a range of other group behaviours such as nest construction or brood relocation where the advantages of communication are less apparent. For example, many ant species regularly move brood within a nest and between nests to raise offspring under optimal temperature and humidity^16,17,18,19,20^. Such controlled responses to environmental variables are a central part of colony organisation in social insects because they have direct impacts on colony growth, metabolic expenditure, survival and reproduction^19,20,21^.

In this study we conduct a detailed analysis of brood transport in the ant *Camponotus fellah* to investigate to what extent workers communicate to displace the brood after changes in environmental conditions. We took advantage of the fortuitous observation that workers moved the brood in response to environmental changes in three colonies (colony size=197, 192, and 206 workers, brood items=150, 60 and 35) to investigate whether workers communicate about observed changes in local conditions. In *C. fellah*, as in most other ants, workers quickly respond to environmental changes to move the brood to the nest regions with the best conditions ^22, 23, 24, 25^.

## Results

### Colonies transport brood in synchrony

In each of the three colonies, and each of the nights, workers responded to the environmental change, initiating brood transport 22.4±6.2 minutes (mean±SEM) after the light was turned off in the tunnel (Fig. 1). There were neither consistent differences across colonies, nor a change in the response delay over the three days (ANCOVA, colony: F=0.9, p=0.37; day: F=0.77, p=0.41; interaction colony*day: F=0.41, p=0.69). On average workers took 160.0±48.0 minutes to move all the brood from the nest to the tunnel once transport was initiated. Workers also performed this task in synchrony with multiple workers transporting in parallel during 66.1±28.0% of the time. The average time taken by a worker to transport one brood item was 36.7±4.0 seconds (see Supplementary Video 1). Workers that transported more brood items were faster to transport brood than those transporting fewer brood items (Spearman rank correlation: *ϱ*=-0.51, p<0.0001; Supplementary Fig. 2). There were again neither significant differences across colonies, nor over days, in the time required to transport all the brood (ANCOVA on log-transformed duration: colony: F=1.5, p=0.31; day: F=1.3, p=0.24; colony*day: F=1.2, p=0.40).

**Fig 1.**
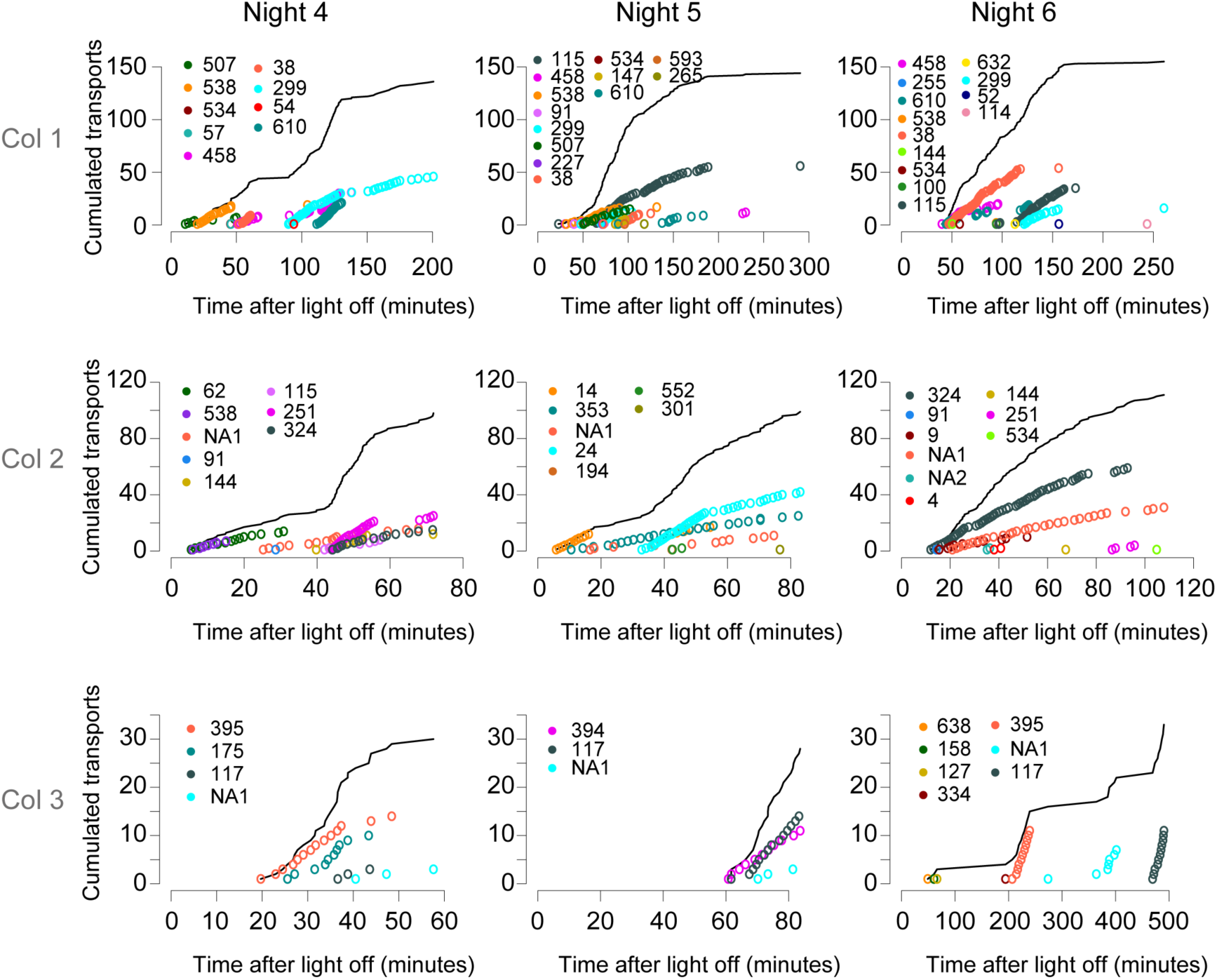
Brood transport dynamics on three consecutive days in three colonies. The black line indicates the cumulated number of brood transports to the tunnel of all workers. Each coloured circle represents a single brood transport event by one worker, and data are shown as cumulative transports. Different colours represent different workers.

### A small minority of a colony’s workforce transports brood

The number of workers involved in brood transport was consistently low, with only 8.1±1.1 workers (4.1%±0.6% of the workforce) participating in brood transport on any given day in any given colony (Fig. 2). Colonies did not differ in the distribution of the workload among workers, and there was no significant change over days in the way the workload was distributed among transporters (ANCOVA: colony: F=0.40 p=0.67; day: F=0.15 p=0.86; colony*day: F=0.14, p=0.97). However, there was variation among transporters in their relative contribution with the notable effect that more than 80% of all brood transports were performed by less than 1.8% of all workers. In addition, there was also a high worker turnover with 66.9±5.2% of the transporters working on a single night, while only 18.8±11.9% of the transporters worked on all three nights. Importantly, however, the persistent transporters were responsible for 44.3±25.3% of all transports while those that worked a single night contributed together to 24.8±18.7% of the transports.

**Fig 2.**
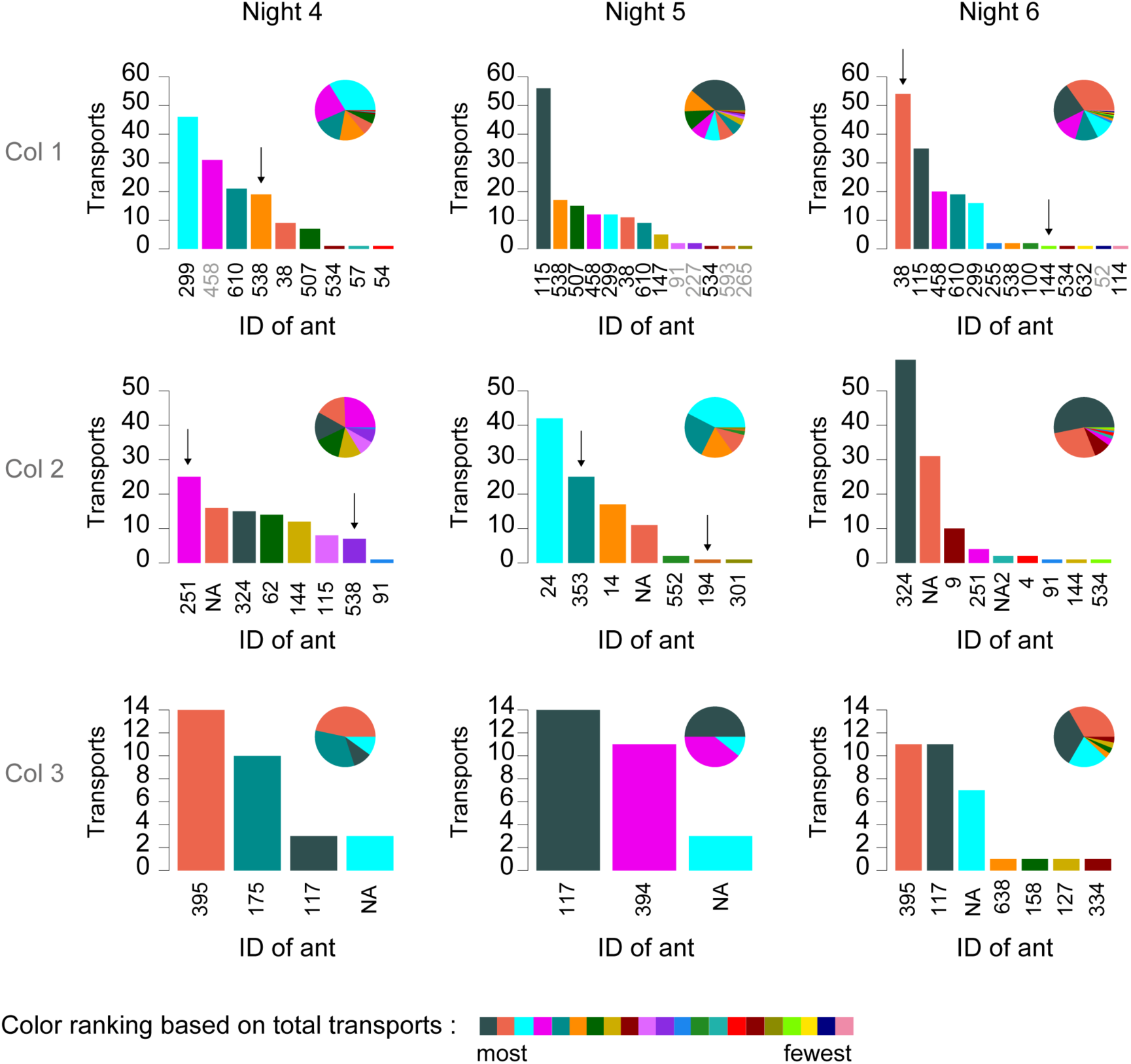
The workload is distributed unevenly among the transporters. Absolute numbers are given in the histogram, and proportions are indicated in the pie chart. Arrows indicate workers that transport without being privately informed (i.e. they had not visited the tunnel before starting to transport). Transporters with ID labels in black are nurses, while those with labels in grey belong to the cleaner or forager groups.

### Transporters are nurses

To determine whether brood transport was preferentially conducted by a specific group of workers, we used the Infomap algorithm^26^ to determine the daily interaction networks of workers and assign each of them to a specific social group^27^. Colonies had on average 55.9%±11.3% nurses, 16.5%±4.9% cleaners and 25.1%±7.4% foragers (Fig. 3). Nurses were 3.8 times more likely to transport than cleaners, and 7.3 more likely to transport than foragers (ANOVA, F=51.38, p<0.0002). There was also an effect of age, with transporters being on average younger (83.5 days) than non-transporters (119.5 days; Kruskal-Wallis: χ^2^=12.1, p<0.001). This effect was due to age differences between the three groups of workers (average age nurses 93.8 days, cleaners 124.2 days, foragers, 159.4 days; Kruskal-Wallis: χ^2^=138.6, p<0.00001). When only nurses were considered, there was no significant age difference between transporters and non-transporters (Kruskal-Wallis: χ^2^=0.81, p=0.37; insufficient data was available to conduct similar tests for nest cleaners and foragers).

**Fig 3.**
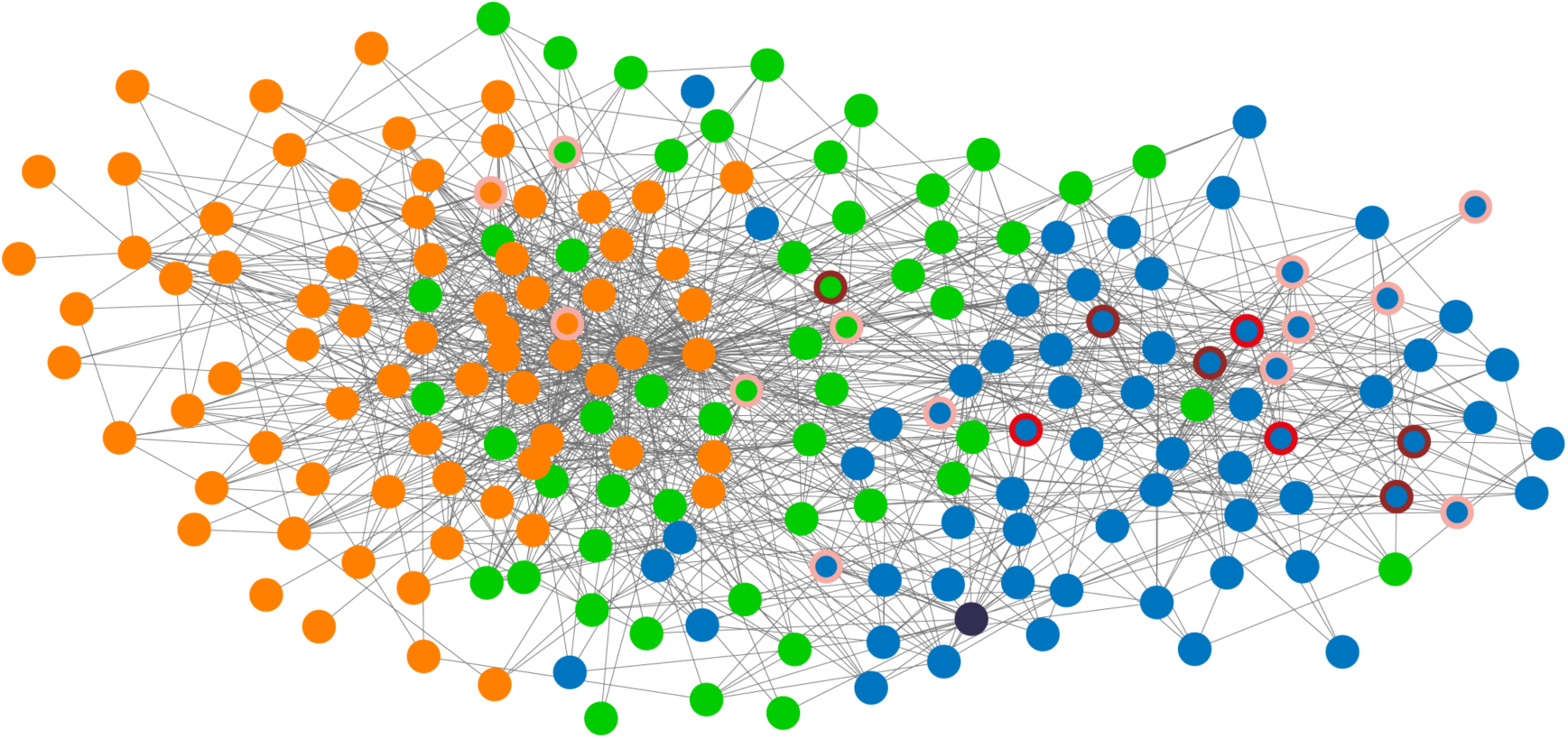
Transporters are mainly nurses. The network shown is that of colony 10 on day 4. Each node represents a worker, and links between nodes are shown for workers who had more than 10 interactions on that day. The network layout is a spring embedded layout. Group membership is indicated by the node colour: nurse (blue), cleaner (green), forager (orange). Red-shaded circles around nodes highlight transporters, with light red indicating transports on one day, medium red indicating transports on two days, and dark red indicating transports on three days.

### Transporters gather information themselves

To determine whether workers make use of information available to others to decide when to initiate brood transport, we tracked the information available to each worker after the light was turned off. Because the nest entrance was constructed with two 90° bends and painted in matt black on the inside thereby preventing light from entering the nest, the only means for workers to know whether there was light in the tunnel was to access it. Workers were therefore considered as having private information once they had left the nest for at least three seconds, which is the minimum amount of time an ant needs to reach the tunnel and return to the nest. Ants were considered as socially informed once they interacted with a privately informed worker.

At the start of brood transport, only 31.6%±2.9% of all workers and 37.8%±8.7% of the nurses had private information. However, almost all transports (99.8%) were performed by privately informed ants. Of the seven workers, which had not visited the tunnel before initiating brood transport, four had transported brood on previous days (Fig. 2). The three remaining workers had visited the tunnel the nights before when it contained brood. Thus, these transporters may have used this information together with circadian timing to initiate transport^24,25^. Therefore, these observations suggest that private information is the primary or only source of information workers use to decide when and where to transport the brood.

### Transporters neither communicate nor recruit help

Five lines of evidence further support the view that workers do not use information obtained from other workers to initiate brood transport. First, transporters did not increase their interaction frequency with other workers once it was dark in the tunnel. The rate of interactions in the hour preceding light-off was not significantly different from the rate during the interval between light-off and the first brood transport (Kruskal-Wallis: χ^2^=0.05, p=0.82; Supplementary Fig. 3). Second, transporters did not change their activity after interacting with a privately informed ant. Their increase in speed — a signature of information transfer in ants^28^ — was similar after interacting with a privately informed or an uninformed ant (Kruskal-Wallis: χ^2^=2.8, p=0.09, see Supplementary Table 1). Third, brood accumulating in the tunnel did not speed up the recruitment of additional transporters. The average time elapsed before one additional worker contributed to brood transport was 16.6±3.4 min. The number of workers already participating in brood transport did not alter the time needed to rally an additional worker (Spearman rank correlation: *ϱ*=0.06, p=0.60; Supplementary Fig. 4). Fourth, the first interaction with a privately informed ant did not trigger a change in behaviour. After interacting with a privately informed ant, transporters and non-transporters were neither more likely to approach the nest entrance (Wilcoxon signed rank test: transporters: V=1232, p=0.79; non-transporters: V=495789, p=0.97) nor to orient towards it (Rao’s spacing test for uniformity: transporters: Test Statistic=139.98, p>0.05 with a critical value=148.34; for non-transporters: Test Statistic=134.13, p>0.05 with a critical value=136.94; Fig. 4A, 4B). Simulations were conducted to determine the expected effect if 90%, 50%, 10% or 0% of the transporters were able to understand a message that they should go to the tunnel after interacting with a privately informed ant (Fig 4C). These simulations revealed that the observed pattern was consistent with a complete lack of communication between privately informed ants and non-informed transporters. Finally, and most importantly we did not observe any successful recruitment through tandem running although these ants are capable of tandem running (see Supplementary Videos 2, 3).

**Fig 4.**
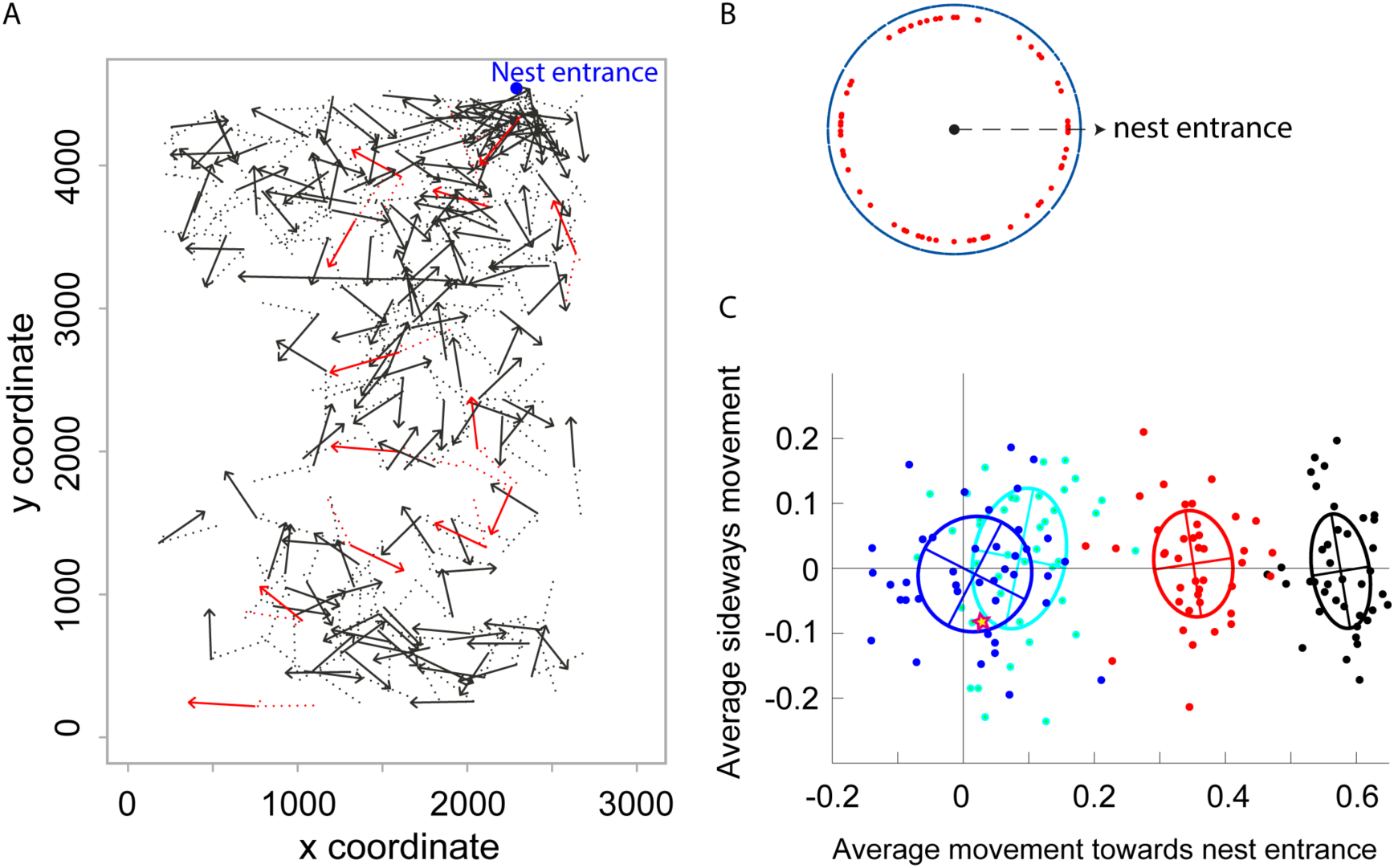
No evidence for communication between workers. (A) Changes of trajectory following the first interaction with a privately informed ant. Arrows indicate the trajectories after the first interaction with a privately informed ant and the dotted lines the trajectories just before this interaction. Transporter trajectories are in red and those of other ants in black. The blue circle indicates the nest entrance. Data shown are those of colony 1 on day 5. (B) Distribution of directions after the first interaction with a privately informed ant. Each dot represents the direction relative to the nest entrance of a single worker on a given day. Red dots indicate transporters and blue dots (forming a ring) indicate other ants. The arrow indicates the direction of the nest entrance. (C) Expected change in direction from simulated data in which 0% (blue), 10% (cyan), 50% (red) or 90% (black) of the ants understood a message. Each dot is the average movement towards the nest entrance of 66 simulated transporters. The cross and ellipse show the average and the standard deviation across 40 simulations with the same set of parameters. The star shows the average of the observed data.

### Colonies do not use quorum sensing to initiate brood transport

At the colony level there was also no indication of a system of quorum sensing leading to the onset of brood transport. At the time of first transport, the percentage of privately and socially informed workers and the percentage of workers in the tunnel varied greatly (privately informed: 0.6% to 12.0%; socially informed: 1.9% to 47.5%, ants in tunnel: 6.0% to 19.4%; Fig. 5). Furthermore, the use of a quorum would imply that colonies deferred the onset of brood transport on some days for almost one hour after reaching the quorum, while starting to transport just minutes after reaching the quorum on other days (delays for privately informed: 4.3–59.8 minutes; socially informed: 2.8–58.8 minutes; ants in tunnel: 5.4–59.1 minutes). Given that the variability was large for both the quorum threshold and the delay until transport onset, it seems unlikely that a minimum colony level information threshold or a minimum ant proportion in the tunnel needs to be reached for brood transport to be initiated.

**Fig 5.**
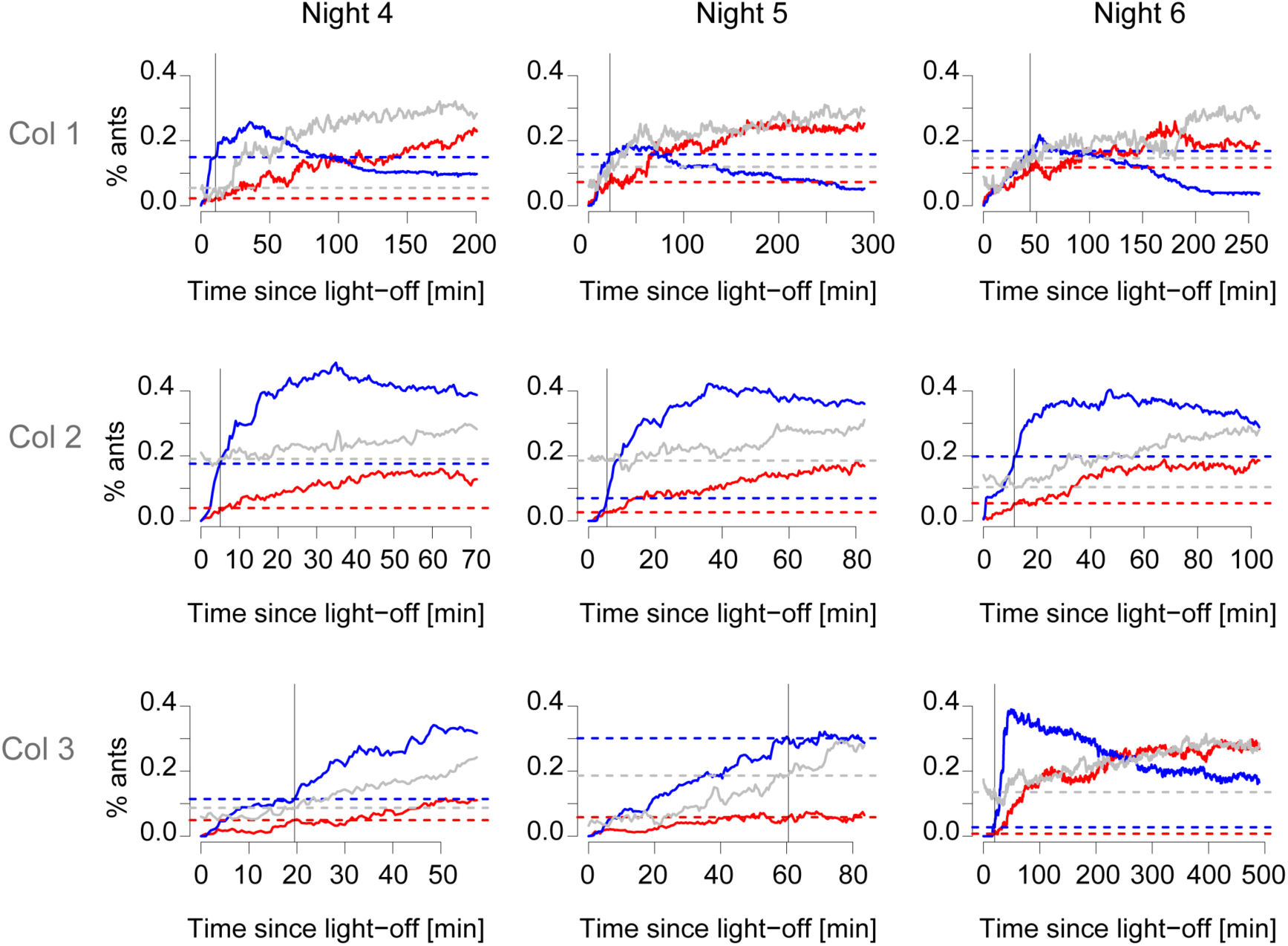
No evidence for a quorum threshold triggering brood transport. Each line shows the percentage of ants: privately informed ants in red, socially informed ants in blue, ants in the tunnel in grey. The vertical line indicates when the first transport occurred, and the dashed lines highlight the percentages of ants at the time of first transport.

Finally, our analyses also revealed high consistency in the direction of brood transport (Supplementary Fig. 5). Overall, there were only 20 return-transports (2.3%) among the 859 transports recorded. Interestingly, the majority of the workers (69.2%) performing return transports did not transport brood to the tunnel while the vast majority (91.7%) of the workers transporting brood to the tunnel did not perform return-transports.

## Discussion

The use of an automated system allowed us to obtain detailed and individual-level information on the processes regulating brood transport in response to environmental changes, a process central to the organization of social insect colonies. Overall, workers quickly transported the brood to the preferable location after the light was turned off, and workers almost never transported brood in the wrong direction. However, this seemingly coordinated transport occurred without any detectable sign of communication among workers. While workers frequently interacted, these interactions resulted in no visible change in the behaviour of the transporters, even if the interaction partner had knowledge about the tunnel being dark. Instead, transporters appeared to rely exclusively on self-gathered information, because they initiated brood transport only after having noticed the change of state of the tunnel themselves. Together, these data indicate that synchronised behaviour at the colony level can occur without communication.

Visual inspections of our videos also revealed no evidence that workers relied on chemical signals to initiate and communicate brood transport. Transporters never dragged their gaster over the ground, as ants typically do when depositing trails. There were also no instances of worker tandem running, thereby excluding targeted recruitment that could have been mediated by secretions from a gland^13^. The only targeted recruitment that we observed was that of the queen and in one instance that of non-transporting workers (see Supplementary Videos 2, 3). In these cases a worker approached the head of the queen or worker and pulled on her mandibles, with the effect that the pulled ant became active and followed the worker in a tandem-run to the tunnel.

The observed lack of communication is likely due to the inherent difficulty of reliably communicating a message in a noisy environment. Communication requires that an informed individual intentionally encodes a message, transmits it successfully, and that an uninformed individual is able to receive it, decode it, and act upon it^29^. Ants have a limited ability to convey a message through tactile communication alone^28,30,31^. In addition, the density of workers is extremely high in the nest, resulting in numerous interactions not only with informed individuals but also with uninformed ones. Such a situation leads to a very noisy system where conflicting feedbacks may readily compromise any attempts of communication. Moreover, investing time in recruiting a helper would only beneficial if the time needed for successful recruitment is short, and if recruitment occurs early on (see Figure 6).

**Figure 6.**
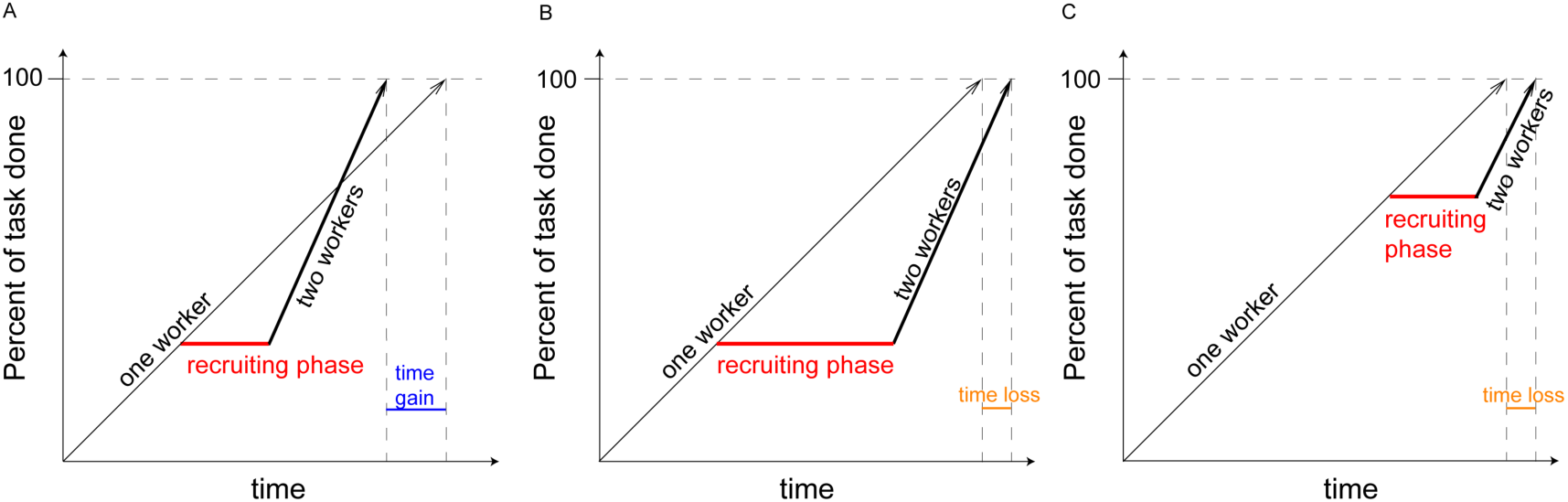
Cost and benefit of successful recruitment. The time invested in recruiting help is indicated in red. The time gained from recruiting a helper is shown in blue, and the time lost due to recruiting help in orange. (A) Recruiting a helper early on after the task is initiated and with little time investment enables faster completion of the brood transport than without a helper. (B) Recruiting a helper early on but with high time investment delays the completion of the brood transport compared to a situation without a helper. (C) Recruiting a helper later while the task is performed also delays the completion of the brood transport compared to a situation without a helper.

Our observation that transporters check the state of the tunnel themselves, before starting to transport brood, suggests that individual workers gather cues from the environment before deciding to transport brood. The most likely cues used by the transporters in our experiments are the confinement, absence of light and presence of workers in the tunnel^32,33^. The use of cues for decision-making also occurs in other ants, and for processes unrelated to brood transport. For instance, in harvester ants, potential foragers decide whether or not to initiate a foraging trip based on the frequency with which they meet returning foragers^34,35^. Workers of the black garden ant *Lasius niger* use the chemical profile of the nest wall and their own body size compared to the height of nest pillars as cues to decide whether to switch from wall building to building a roof^36^. These data, together with our results, suggest that the use of cues as a mean to obtain private information might be more widespread and easier to implement in ant colonies than information exchange through tactile communication.

The use of cues combined with the lack of communication and the absence of a quorum means that transporters most likely decide independently of each other whether, when and where to transport the brood. Such individual-led decisions are further supported by rare instances in which a worker mistakenly returned brood from the tunnel to the nest, while transporters were already moving brood to the tunnel. Interestingly, the vast majority of transporters arrived at the same decision and transported brood from the nest to the tunnel. This strong uniformity in behaviour suggests that there is high homogeneity in preferences among group members.

Our results indicate that colonies can display synchronized behaviour without communicating thus emphasizing that not all group-level behaviours in social insects are driven by communication. We suspect that communication is context-dependant and only used when cue-based options are insufficient. For instance, the communication that precedes brood transport in house-hunting ants occurs in the context of an emergency after their nest has been destroyed^9,14^. In contrast, synchronization without communication is optimal when reliable communication is expensive, hard to achieve, or when perfect synchrony is not needed^29,37^. It can be achieved if workers share similar preferences and react to the same cues, which are limited in time. In our experiments light in the tunnel acted as this strong time-limited cue. Synchronized group behaviour exists also in solitary bees, who congregate at nesting sites for reproduction^38^, bats and starlings that converge at seasonal feeding and sleeping spots^39,40^ and Mormon crickets, who migrate in masses in search for salt and proteins^41^. In ants simulations further suggest that food choice during foraging could be achieved without communication through individual learning and preference^42^.

Our results also revealed that only a tiny fraction of the individuals, 1.5%–6.6% of the colonies’ workforce —as few as three workers in some cases— contributed to brood transport. Moreover, within colonies there was strong variation in the relative contribution of workers with more than 80% of all transports being carried out by less than 1.8% of the workers. Similar fractions of transporters and workload disparities were observed in colony emigrations of *Formica sanguinea* and *Camponotus sericeus^43^*. The large variability in behaviour is puzzling and we offer two possible explanations. There could be specialist nurses that focus on brood transport. Indeed nine out of 48 transporters moved brood every single night and did slightly less than half of the work, thus acting as key individuals^44^ during the brood displacement. Similar specialization has been reported for foraging, brood care, stone collection^45,46,47^ and could result from inherent and consistent differences between workers, for example in motivation, physiology, or sensory threshold^48,49^. Another explanation is that transporters represent a varying subset of the nurses, whose likelihood to transport depends on the individual’s state in the early night. This idea is supported by the observation that two thirds of the transporters only worked a single night.

Importantly, a small minority of transporters imposed their transport decision on the colony. Such an outcome was only possible because the other workers did not oppose the brood transports or if they did so initially, never persisted in their opposition. Minority-driven behaviour occurs also in *Paratrechina longicornis* ants, where a single worker can temporarily decide the pull direction during collective transport^50^. Our results therefore highlight that a small minority of the workforce can determine the colony fate through persistent activity in a largely indifferent society. Similar observations exist for fish schools and human crowds where few knowledgeable individuals can lead large groups of uninformed individuals to a new location^51,52^. Ultimately, the social unresponsiveness of the majority might be the optimal strategy because social unresponsiveness can ensure that the colonies react to environmental change while also being robust to noise and avoiding losses in information accuracy resulting from an over-reliance on social information53.

## Acknowledgements

We thank M. Chapuisat, O. Feinerman, N. Stroeymeyt, T. Richardson, T. Kay and S. McGregor for comments on an earlier version, and B. Sutcliffe for proofreading. DPM was supported by a grant from HFSP, JPE was supported by an advanced ERC grant, LK was supported by grants from the Swiss NSF and an advanced ERC grant.

## Contribution

DPM and LK planned the experiment. DPM and AC designed the experimental system and performed the experiment. DPM and JPE analysed and interpreted the data. DPM wrote the paper with input from JPE and LK. All authors revised the paper.

## Material and Methods

The three colonies were each established from a single queen collected after a mating flight in Tel Aviv on March 23^rd^ 2007. The experiment started when queens were 3 years old, out of a maximum life span of 26 years^54^. At the start of the experiment, colonies each comprised a queen, brood and 197, 192 and 206 workers, for colonies 1, 2 and 3 respectively. The colony sizes were those naturally reached by queens of that age, and reflect normal growth rates in the laboratory; no data are available for field colonies. All workers were the offspring of a single queen, which in *Camponotus fellah* is usually singly-mated^55^.

To determine workers’ age, new-born workers were paint-marked on a weekly basis during the 12 months preceding the experiment. Because 38 out of the 45 transporters were nurses, we limited the analysis on the effect of age to nurses only.

During experiments colonies were kept in a dark nest chamber connected by a 60 cm long and 1cm wide tunnel to a foraging chamber. The tunnel and the foraging box had 12h light-12h dark cycles, and the ants had access to food (gelatinous sugary water) and water in the foraging box. The temperature (30 °C), humidity (60%), light (~500 Lux), and food supply were computer-controlled, and both chambers were filmed from above with high-resolution monochrome cameras operating under infrared light, as previously described^27^ (Supplementary Fig. 1). All colony members were video-tracked using fiducial identification labels over 14 consecutive days. We recorded the position and orientation of all individuals twice per second.

During the night, workers transported the brood to the tunnel and brought it back to the nest at dawn, presumably because they prefer to keep the brood in a confined environment rather than an open environment when both are dark. We tracked the transport of brood items manually during three consecutive nights. A brood transport was defined as the time interval from when an ant collected one (or several) brood items from the nest box, to when the ant disappeared with it into the tunnel. We also recorded cases where brood was transported from the tunnel to the nest. In these return-transports, the transport was defined as the time interval from when the ant entered the nest with brood until the ant dropped the brood. For each transporter and each night we defined its workload as the number of transports during that night and its work time as the time from the start of its first transport until the end of its last transport. Using the work times of all workers, we estimated synchrony as the percentage of time during which at least two workers worked in parallel. We also visually inspected the videos for instances of tandem running, that is events where one ant guides another ant to the tunnel. A tandem-run results in successful recruitment if the follower ant subsequently starts transporting brood. We did not track brood transports in the mornings when the lights turned on in the tunnel, because in these conditions all ants in the tunnel were immediately informed of the environmental change, thus making the question of communication inane.

To determine group membership of each worker, *i.e*. nurse, cleaner or forager, we used the same approach as in Mersch *et al*. (2013)^27^. In brief, we inferred all social interactions between workers based on their distance and orientation, and analysed the social networks with the Infomap algorithm^26^ to assign each worker to a group. Because the majority of workers were in the tunnel at night and thus undetectable with our tracking setup, we built daily interaction networks using only data collected between 8am and 7pm, when the majority of workers were detectable.

To measure the speed change following interactions, we calculated the speed during the 10 seconds prior to the interaction and during the 10 seconds after the interaction. We included only those interactions for which we had data on the speed before the interaction for both partners and on the speed after the interaction for the focal ant. As a consequence, 50 interactions (10.2%) were excluded from the analysis. Excluding these interactions had neither an impact on the average duration of an interaction (10.5±29.9 s *vs*. 10.3±30.5 s) nor on the proportion of interactions with privately informed partners (7.72% *vs*. 7.69%). To further ensure that our results are not influenced by the chosen interval (10 s), we repeated the same analyses for shorter (5 s) and longer (20 s) time intervals. Because the results were the same for all time intervals (see Supplementary Table 1), we only report data for the 10-second interval.

To investigate whether a privately informed ant can communicate information about the change of state in the tunnel to its interaction partner we estimated the change in trajectory of each worker following its first interaction with a privately informed ant. We calculated the heading of the ant’s trajectory after it had moved away from the interaction point, transforming data of all colonies so that an orientation of 0° corresponds to an orientation towards the nest entrance. We also calculated the distance to the entrance at the time of the interaction and after the ant had moved at least 2 cm (≈ queen body length) away from the interaction point. Workers who did not interact with a privately informed ant before the end of the brood transport were not included in the analysis (351 out of 1785 ant-days excluded).

To estimate how communication about the change of state in the tunnel could modify the trajectory of workers, we generated simulated datasets in which 0%, 10%, 50% or 90% of the transporters moved toward the nest entrance after interacting with a privately informed ant. Understanding the message meant that one bit —that is, one unit of information— was transferred from the privately informed ant to the transporter. Such one-bit information could convey two options —towards and away from nest entrance— and signal to the transporter to move towards the nest entrance. Each dataset was the average of 66 simulated direction vectors v_j_ defined as

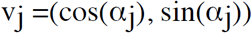

with α_j_ being the angle of the direction relative to the line connecting the interaction point with the nest entrance. For each informed transporter, we randomly chose a direction from a uniform distribution limited to angles between –π/2 and π/2, for all other transporters we randomly chose an angle from a uniform distribution between – π and π. We repeated this process 40 times for each information level. We also calculated the average direction of the 66 transporters from the observed data.

To test whether a quorum triggered the observed brood transport, we determined the number of ants, the number of informed ants, and the number of ants in the tunnel at the time of the first brood transport. To estimate whether the quorum induced brood transport, we also calculated the duration between the time the quorum was reached for the first time and the first brood transport. Because the estimated quorum varied between colonies and days, we calculated the delays for all colonies and days using the smallest estimated quorum threshold.

We performed all statistical analysis in R (Version 3.4.0)^56^. When the test assumptions were met, we used two-tailed parametric tests and included the colony ID as a random factor in our analysis; otherwise we used non-parametric tests. For statistical tests on colonies, each colony was one replicate. For statistical tests on individual workers, each transporter on each day was a replicate. The data analysis code will be available as a zip file.

The data used to prepare all figures and perform statistical tests will be available on Dryad DOI after publication in a journal.

## Supplementary material

**Supplementary Figure 1:**
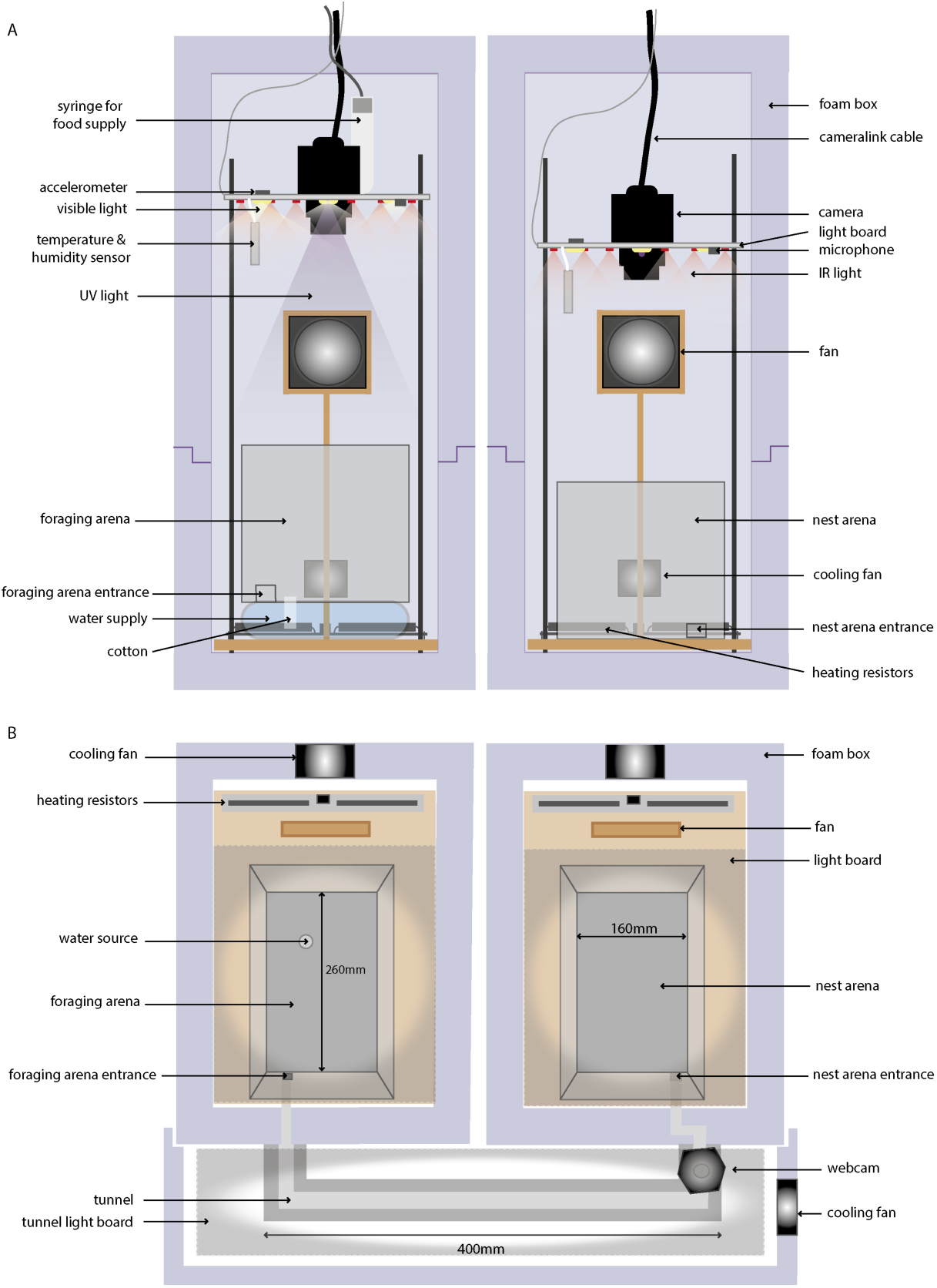
Tracking setup (A) Lateral view (B) Top view; reproduced with permission from Mersch *et al*. (2013)^27^

**Supplementary Figure 2.**
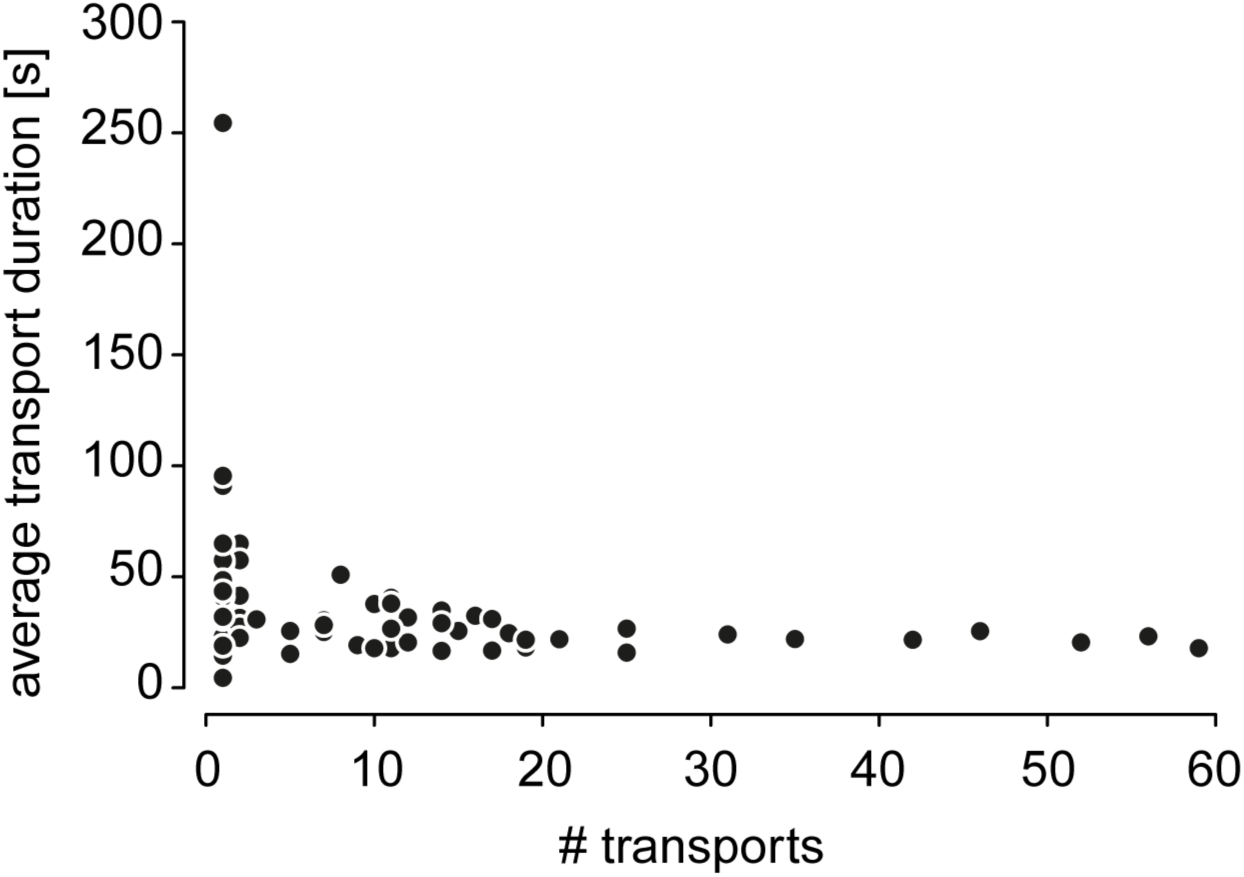
Individual workers transport brood rapidly. Each black dot shows the average transport time needed by a single transporter.

**Supplementary Figure 3.**
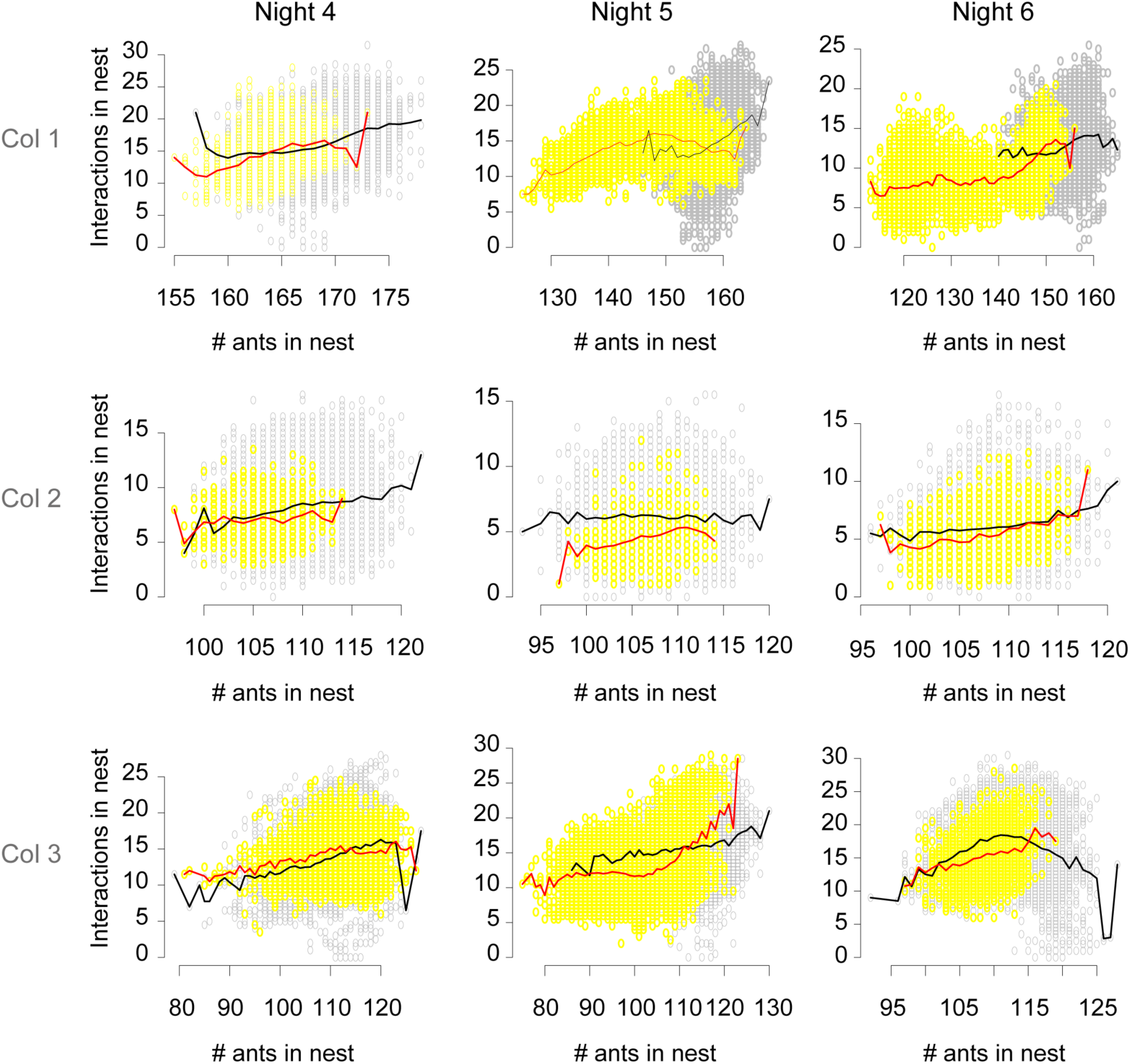
No change in interaction frequencies after light-off. Grey dots show data in the hour preceding light-off. Yellow dots show data between light-off and the first transport. The black line shows the average relationship between the number of ants in the nest and the number of interactions before light-off, and the red line shows the same relationship in the interval between light-off and the first brood transport.

**Supplementary Figure 4.**
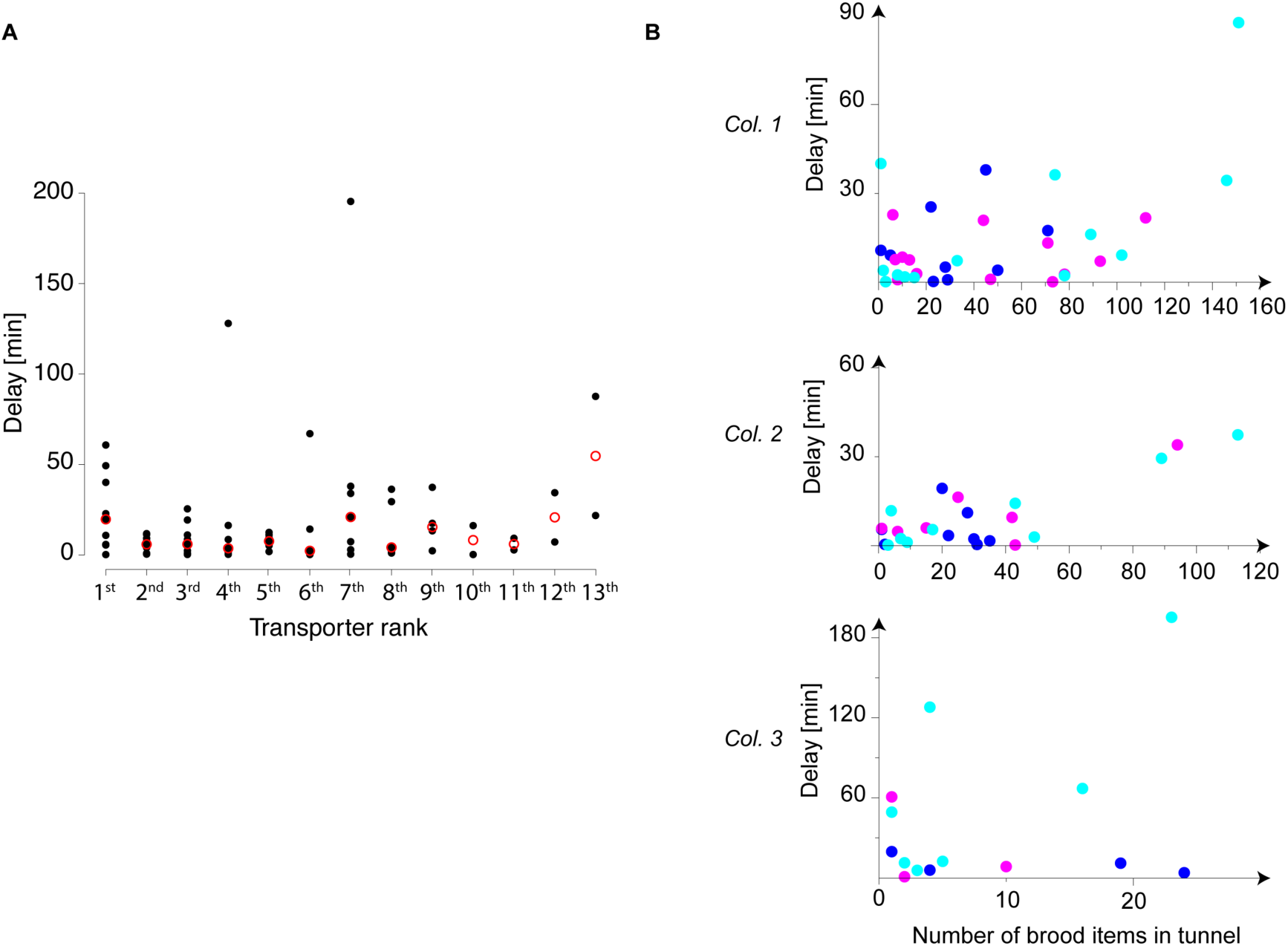
Brood accumulation in the tunnel does not speed up transporter recruitment. A. Each black dot shows the recruitment delay. For all but the first transporter, recruitment delays are with regard to the transport start of the previous transporter. For the first transporter, recruitment delays are with regard to light-off. Red circles indicate the median recruitment delay for each transporter rank. B. The recruitment delays are the same as in A. Blues dots show data for night 4, magenta dots data for night 5, and cyan dots data for night 6. Data are shown separately for each colony.

**Supplementary Figure 5.**
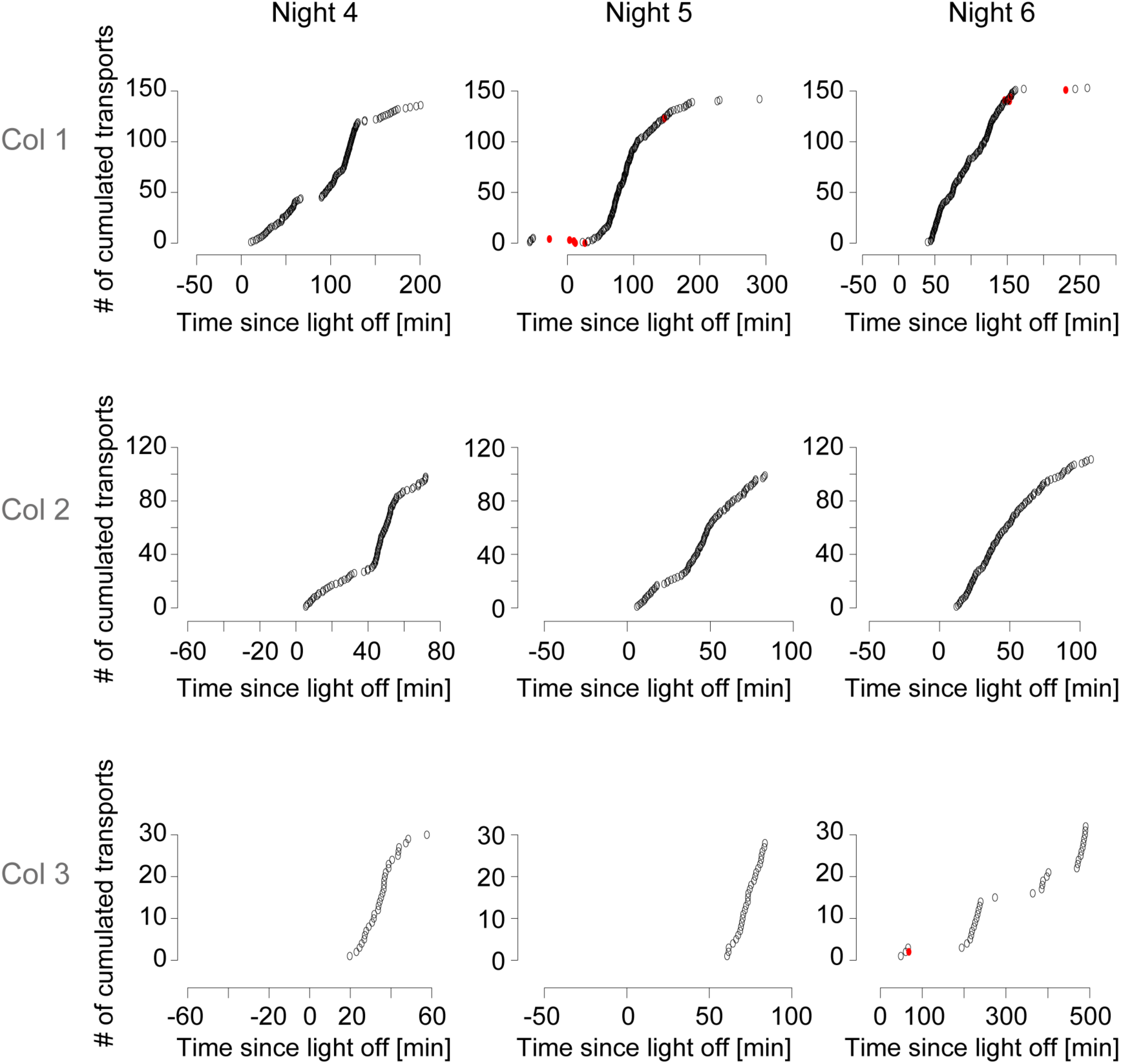
Workers transport almost exclusively from the nest to the tunnel. Grey dots show transports from the nest to the tunnel. Red dots show transports from the tunnel to the nest.

**Supplementary Table 1.**
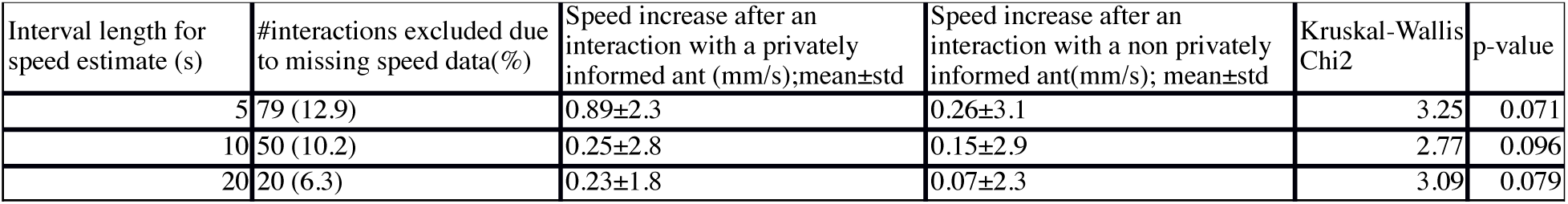
Speed change after an interaction with a privately informed ant.

**Supplementary Video 1. Worker transporting brood.** Worker 62 transports brood to the tunnel. At 16s in the video, ant 62 takes brood directly from another worker without this worker changing its behaviour. Data is from colony 2 and the frame rate is accelerated 5 times. The green line shows the worker’s trajectory in the previous minute.

**Supplementary Video 2. Targeted queen recruitment to the tunnel.** Worker 632 (in pink) approaches the queen, pulls on her mandibles, and then returns to the tunnel with the queen (in blue) following her. The data are from colony 1.

**Supplementary Video 3. Recruitment of two non-transporters to the tunnel.** Worker 458 (in green) interacts with workers 607 (in blue) and 278 (in cream), and both then follow worker 458 to the tunnel. The trajectories are shown for all three workers after the interactions finished. The data are from colony 1.

